# Structural studies of MMP-3 interaction with triple-helical collagen introduce the enzyme’s new roles in tissue remodelling

**DOI:** 10.1101/756460

**Authors:** Szymon W. Manka, Dominique Bihan, Richard W. Farndale

**Author notes:** MRC Prion Unit, UCL Institute of Prion Diseases, 33 Cleveland Street, London, W1W 7FF, UK.

## Abstract

Matrix metalloproteinase-3 (MMP-3 or stromelysin 1) participates in normal extracellular matrix (ECM) turnover during embryonic development, organ morphogenesis and wound healing, and in tissue-destructive diseases, such as aneurysm, cancer, arthritis and heart failure. Despite its ability to hydrolyse numerous proteins in the ECM, MMP-3 fails to cleave the triple helix of interstitial fibrillar collagens. Nonetheless, it can still bind to these collagens although the mechanism, location and role of binding are not known. We used the Collagen Toolkits, libraries of triple-helical peptides that embrace the entire helical domains of collagens II and III, to map MMP-3 interaction sites. The enzyme recognises five sites on collagen II and three sites on collagen III. They share a glycine-phenylalanine-hydroxyproline/alanine (GFO/A) motif that is recognised by the enzyme in a context-dependent manner. Neither MMP-3 zymogen (proMMP-3) nor the individual catalytic (Cat) and hemopexin (Hpx) domains of MMP-3 interact with the peptides, revealing cooperative binding of both domains to the triple helix. The Toolkit binding data combined with molecular modelling enabled us to deduce the putative collagen-binding mode of MMP-3, where all three collagen chains make contacts with the enzyme in the valley running across both Cat and Hpx domains. The observed binding pattern casts light on how MMP-3 could regulate collagen turnover and compete with various collagen-binding proteins regulating cell adhesion and proliferation.

## Introduction

Fibrillar collagens I, II and III are the major components of the extracellular matrix (ECM). They provide tissues with tensile strength but also constitute active scaffolds, presenting sites for cell adhesion, for attachment of other ECM components and for storage of many regulatory and signalling molecules^1,2^. Each collagen consists of three polypeptide α-chains folded into a unique triple helix, with the characteristic Gly-X-Y repeats (where X and Y are often proline and hydroxyproline, respectively) of these chains offset axially by one residue, resulting in distinct leading, middle and trailing chains^3^. Collagens II and III are homotrimers with identical α-chains, whereas collagen I is a heterotrimer comprised of two α1 chains and one α2 chain. These trimers are called tropocollagens and contain an uninterrupted 300 nm long triple helix flanked by short non-helical extensions (telopeptides). At the next level of collagen structure, five tropocollagens assemble side-by-side as a microfibril, with a 64-67 nm (one D-period) axial stagger. One may envisage such blocks organised head-to-tail with a 0.54D gap between them in the collagen fibre^4,5^, resulting in fibrils and fibres with the canonical D-periodic cross-striation observed using transmission electron microscopy^6,7^.

Collagen turnover is slow during homeostasis, but relatively rapid in development, organ morphogenesis, tissue remodelling and wound healing^8^. It is also dramatically unbalanced in fibrotic or degenerative diseases, such as arthritis, atherosclerosis, aneurysm and cancer^9^. Matrix metalloproteinases (MMPs) play major roles in these physiological and pathological processes as they collectively cleave most, if not all ECM components^10^, and can generate paracrine bioactive products^11,12^. Among them are collagenases: MMP-1, −8 and −13, and stromelysins: MMP-3, −10 and −11, all secreted as inactive zymogens comprising an N-terminal pro-domain and a catalytic domain (Cat) connected to the C-terminal hemopexin domain (Hpx) via a flexible linker (hinge region). The main difference between these two groups is that stromelysins do not cleave the triple-helical regions of fibrillar collagens, despite sharing a similar degree of structural identity amongst themselves and with the collagenases (>50 % sequence identity and virtually superposable tertiary structures of their Cat and Hpx domains^10,13^. Although MMP-3 is not directly collagenolytic, it is a critical procollagenase activator^14,15^. The prototypic collagenase MMP-1 has only 10-20% of its maximal collagenolytic activity without proteolytic activation by MMP-3^14^. Hence, proMMP-3 is often secreted together with proMMP-1 by mesenchymal cells, macrophages and cancer cells stimulated with pro-inflammatory cytokines^12^.

Besides activating procollagenases and other proMMPs MMP-3 itself breaks down: i) multiple ECM components, including proteoglycans, fibronectin, laminin, collagen telopeptides and basal lamina collagen IV, ii) cell surface proteins, such as E-cadherin at cell junctions, and iii) other non-ECM molecules influencing cell proliferation and differentiation^16^. The enzyme is upregulated in many diseases. For example, after myocardial infarction it may serve as a predictor of adverse left ventricular remodelling and dysfunction^17^ and in idiopathic pulmonary fibrosis it may disrupt lung epithelia through cleavage of E-cadherin^18^, causing epithelial-to-mesenchymal transition. MMP-3 is also found in involuting mammary gland, cycling endometrium, long bone growth plate, atherosclerotic plaque, gastrointestinal ulcers, around various tumours and in other tissue remodelling contexts, reviewed by Nagase^16^. MMP-3 is thus considered an upstream regulator of tissue remodelling in health and disease^19^, but its exact *in vivo* roles are not clearly established. For example, in rheumatoid arthritis, MMP-3 is abundantly expressed in cartilage and serum levels are used in diagnostics, but MMP-3 knockouts give conflicting outcomes in different disease models^20–23^.

Although MMP-3 does not cleave triple-helical regions of fibrillar collagens, it can bind to them^24,25^, which may be the basis of its tissue retention and which may directly or indirectly impact collagen and noncollagen matrix turnover. Here, we report the locations of MMP-3 binding sites along tropocollagens II and III using the THP Toolkits that proved useful in mapping MMP-1^26^ and MMP-13^27^ footprints on these tropocollagens and those of many other collagen-binding proteins^28,29^. The Toolkit screening reveals that MMP-3 binding to triple-helical collagen requires Phe residue in position X of the Gly-X-Y repeat and that the recognition of this critical motif relies on cooperative binding of both Cat and Hpx domains. Having combined computational methods with experimental restraints we propose an integrative model of an MMP-3-collagen complex. We show that the multi-site binding of MMP-3 to fibril-forming collagens can influence their fibrillation, which *in vivo* may alter their exposure to collagenases, providing an additional mechanism of regulation of collagen degradation. Lastly, we discuss the potential consequences of MMP-3 binding to collagens II and III with respect to other binding partners of these collagens.

## Results

### Both Cat and Hpx domains of MMP-3 participate in the binding of the triple helix

We synthesised 56 THPs of Toolkit II and 57 THPs of Toolkit III. Every THP in each Toolkit contains 27 amino acids (aa) of the respective collagen (guest) sequence, flanked by 5 GPP repeats and a GPC triplet (host sequence). The first and the last 9 aa of the guest sequence overlap with the preceding and the consecutive THP in the series, respectively. We used a solid phase binding assay to screen these peptide libraries for binding of biotinylated proMMP-3(E200A), mature MMP-3(E200A) and the isolated Cat and Hpx domains. The active site mutation (E200A) does not affect MMP-3 conformation and is used to prevent autolysis and autoactivation. ProMMP-3(E200A) and the Cat and Hpx domains alone showed no prominent binding to any of the Toolkit peptides (Fig. 1A). Only the mature MMP-3(E200A) specifically recognised 9 THPs in Toolkit II: 9, 10, 13, 22, 23, 35, 36, 39 and 45 (A450 ≥ 0.1, except peptide 1, where the signal does not appear specific due to high background), and 5 THPs in Toolkit III: 6, 23, 36, 39, 40 (A450 > 0.1). THP III-44, which contains the canonical collagenase cleavage site, may also be considered to have marginal affinity to MMP-3. Some THPs, like III-36 and III-40, seem to weakly interact with proMMP-3(E200A), but this binding appears negligible compared to the activated MMP-3(E200A) (Fig. 1A).

**Figure 1.**
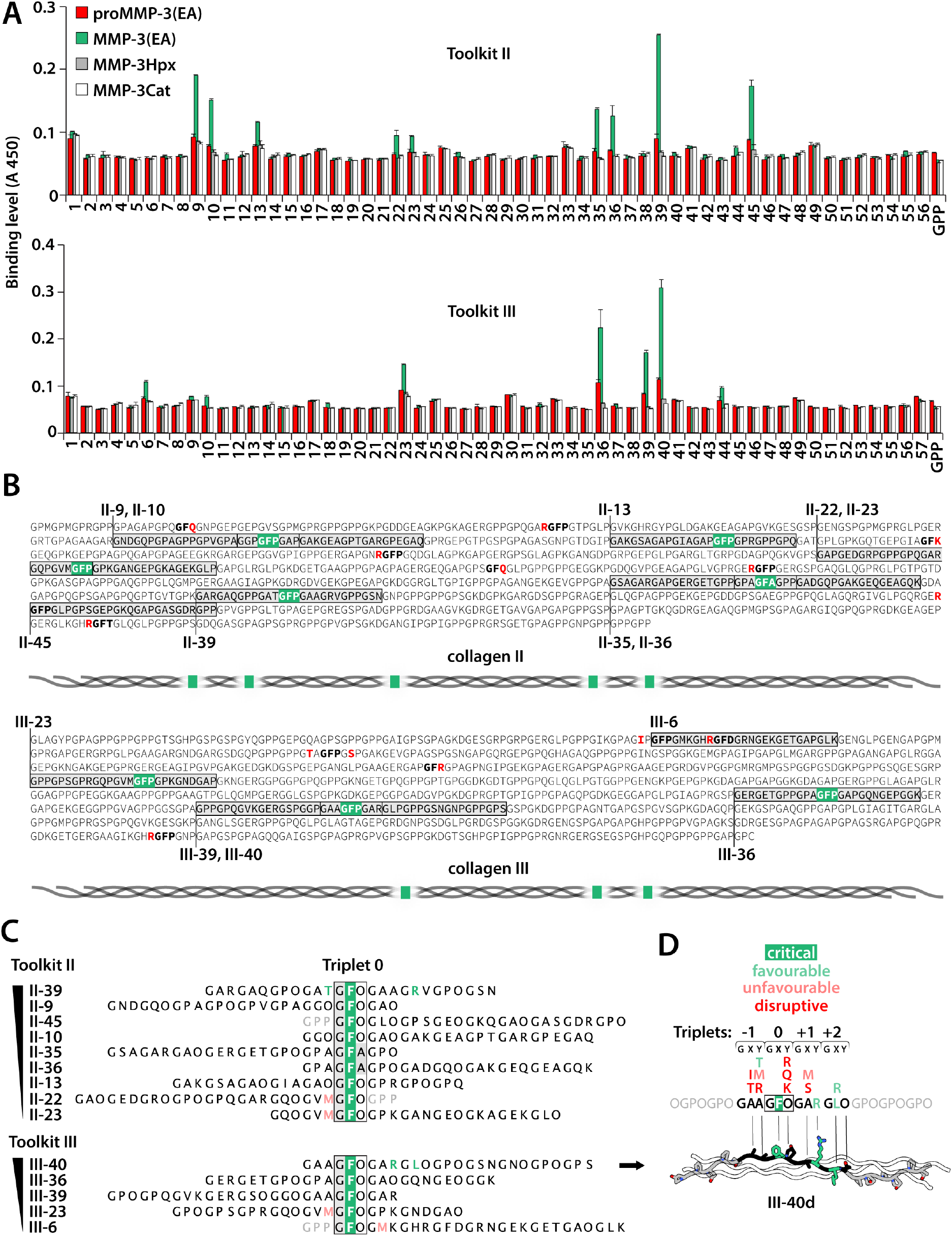
MMP-3 binds 5 sites in collagen II and 3 sites in collagen III via context-dependent recognition of the GFO/A triplet. **A**) Relative binding of biotinylated MMP-3 constructs to Toolkits II and III. The results are the average of 3 independent binding assays performed at room temperature (~20 °C). Error bars, standard deviation (SD). GPP, THP with repeating Gly-Pro-Pro sequence. **B**) Analysis of collagen sequence determinants for MMP-3 binding based on the Toolkit data shown in A. MMP-3-bound peptides are located in the triple-helical domains of human collagens II and III (Uniprot identifiers: P02458 and P02461, respectively). The sequences do not include post-translational modification of proline (P) at the position Y of the GXY repeat to hydroxyproline (O). MMP-3 binds GFO/A (here GFP/A)-containing sites, unless they are preceded or followed by residues marked in red. The critical GFP/A triplet is highlighted in green wherever it mediates MMP-3 binding. Included are schematic overviews of the MMP-3 binding site distributions along tropocollagens II and III. The GFP site represented by peptides II-44/45 is expected to be not recognised by MMP-3. Peptide II-45 is bound only because it starts from the critical GFO triplet and therefore does not contain the preceding R residue, while the overlapping peptide II-44 is not bound. **C**) Sorting the MMP-3-bound peptides according to relative MMP-3 binding affinity. Peptides are aligned on the critical GFO/A triplet designated Triplet 0. One triplet of the *guest* sequence is included (grey font) for peptides starting or ending with Triplet 0. Residues that appear as advantageous for the binding are coloured green and those that appear unfavourable are coloured pale red. **D**) Summary of favourable and unfavourable sequence motifs for MMP-3 binding shown together with a computationally modelled triple-helical peptide (THP) III-40d, designed for modelling of the MMP-3-THP complex. The THP is based on the most optimal MMP-3 recognition sequence determined in peptide III-40 (highest apparent MMP-3 affinity among all Toolkit peptides), consisting of 4 triplets designated: −1, 0, +1, +2. These and the flanking GPO triplets were generated by *mutating* collagen III peptide 1BKV^30^ in *Coot*^31^. For clarity, only one THP chain is shown with coloured residue side chains. Element colours: N, navy blue; O, red.

### MMP-3 recognises triple helices encompassing GFO/A triplets in particular contexts

All the peptides that firmly interacted with MMP-3(E200A) have one unifying feature, the GFO/A triplet, which we designated Triplet 0 (Fig. 1B and C). Notably, Phe at position X is not recognised by MMP-3(E200A) when accompanied by Arg, Gln or Lys at position Y (Fig. 1B and D).

Binding to Triplet 0 depends on a wider sequence context, since it is found at 4 and 3 sites on collagens II and III, respectively (Fig. 1B), that are not recognised by MMP-3(E200A) according to the Toolkit screening. In these cases, the Triplet 0 occurs in the following contexts: R-GFO, IP-GFO or TA-GFO-GS. Therefore, Arg directly preceding the Triplet 0, Ile or Thr at position X of the preceding triplet (Triplet −1) and/or Ser at position X of the following triplet (Triplet +1) appear to disrupt MMP-3(E200A) binding (Fig. 1B and D). This implies that THPs II-45 and III-6 get recognised only because their guest sequences start from the Triplet 0 and, therefore, exclude the disruptive residues that precede them in native collagens (Fig. 1B). This is supported by the clear lack of recognition of the neighbouring THPs II-44 and III-5 that overlap the sites in question (Fig. 1A). We thus conclude that THPs II-45 and III-6 do not represent true binding sites of MMP-3 on native tropocollagens. Consequently, there are 5 and 3 unique interaction sites for MMP-3 on tropocollagens II and III, respectively (Fig. 1B).

To identify the more and less favourable residues for MMP-3 binding in the vicinity of Triplet 0 we ordered the MMP-3(E200A)-binding THPs according to their relative affinity to MMP-3(E200A) (Fig. 1C). Our analysis indicates that Met at position Y of the Triplet −1 or at position X of the Triplet +1 is unfavourable, while Thr at position Y of the Triplet −1, Arg at position Y of the Triplet +1 and Leu or Arg at position X of the Triplet +2 are favourable for MMP-3 binding (Fig. 1C and D).

### Integrative modelling of the MMP-3-THP complex

We predict modes of MMP-3 binding to triple-helical collagen by integrating the established THP binding specificity with the mechanistic insights derived from the Toolkit screening: i) the pro-domain of MMP-3 zymogen interferes with THP binding, ii) both the Cat and Hpx domains of MMP-3 are required for that binding. The sequence of the most strongly bound THP III-40 served to generate a computational THP model III-40d for optimal molecular docking to an MMP-3 model (Fig. 1D). The THP III-40d contains 5 triplets (−1 to +3) of the THP III-40 and 2 flanking GPO repeats built on the backbone of an X-ray THP structure 1BKV.pdb^30^ (Fig. 1D).

To fully explore the experimental restraints for the subsequent docking experiment, we modelled both the pro-form and the mature form of MMP-3, due to the lack of their experimental structures. ProMMP-3 homology modelling was based on proMMP-1 crystal structure (1SU3.pdb^32^), and the full-length mature MMP-3 was based on several homologous structures available: active MMP-1 (2CLT.pdb^33^), MMP-1(E200A) in a complex with a THP (4AUO.pdb^26^) and two structures of MMP-13 (4FU4.pdb and 4FVL.pdb^34^). All models were obtained with Modeller^35^ and were of highest quality, reaching maximal GA341 scores = 1.0 (native-like structures)^36,37^. The best structure in each collection of models was selected according to the lowest Discrete Optimized Protein Energy (DOPE) scores^38^ (see details in Fig. S1).

Since Phe in the Triplet 0 is central to MMP-3 binding, the complex formation must be driven by a hydrophobic interaction. We visualised hydrophobic patches on the surface of the MMP-3 model using hydrophilicity scale proposed by Moon and Fleming^39^, with values ranging from −2.2 for Phe (most hydrophobic) to 5.39 for Lys (most hydrophilic) (Fig. 2). We considered the hydrophobic regions that: i) accommodate a triple helix, ii) involve both Cat and Hpx domains in the resultant THP binding mode, and iii) are sensitive to the presence of the pro-domain. We identified 4 candidate solutions that would satisfy all these experimentally-derived criteria: 2 pertain to collagen binding at the front, and 2 at the back of the enzyme (Fig. 2A). The two frontal binding possibilities (solutions 1 and 2) would be directly blocked by the pro-domain, whereas the two back binding possibilities (solutions 3 and 4) were also considered, since the relative positions of the Cat and Hpx domains could be altered by the pro-domain allosterically and perturb the collagen binding site (Fig. S2). The THP model III-40d was docked to the MMP-3 model in the four proposed ways (Fig. 2A), yielding excellent complexes after Rosetta refinement^40^ in all cases (Fig. 2B and C). Complex 3 is unique in that it engages not only the two domains of MMP-3, but also the linker region via hydrophobic interaction mediated by Leu at position X of the Triplet +2. Each complex bends the triple helix somewhat, especially complex 1. This however does not affect the geometry of the THP (Fig. 2C).

**Figure 2.**
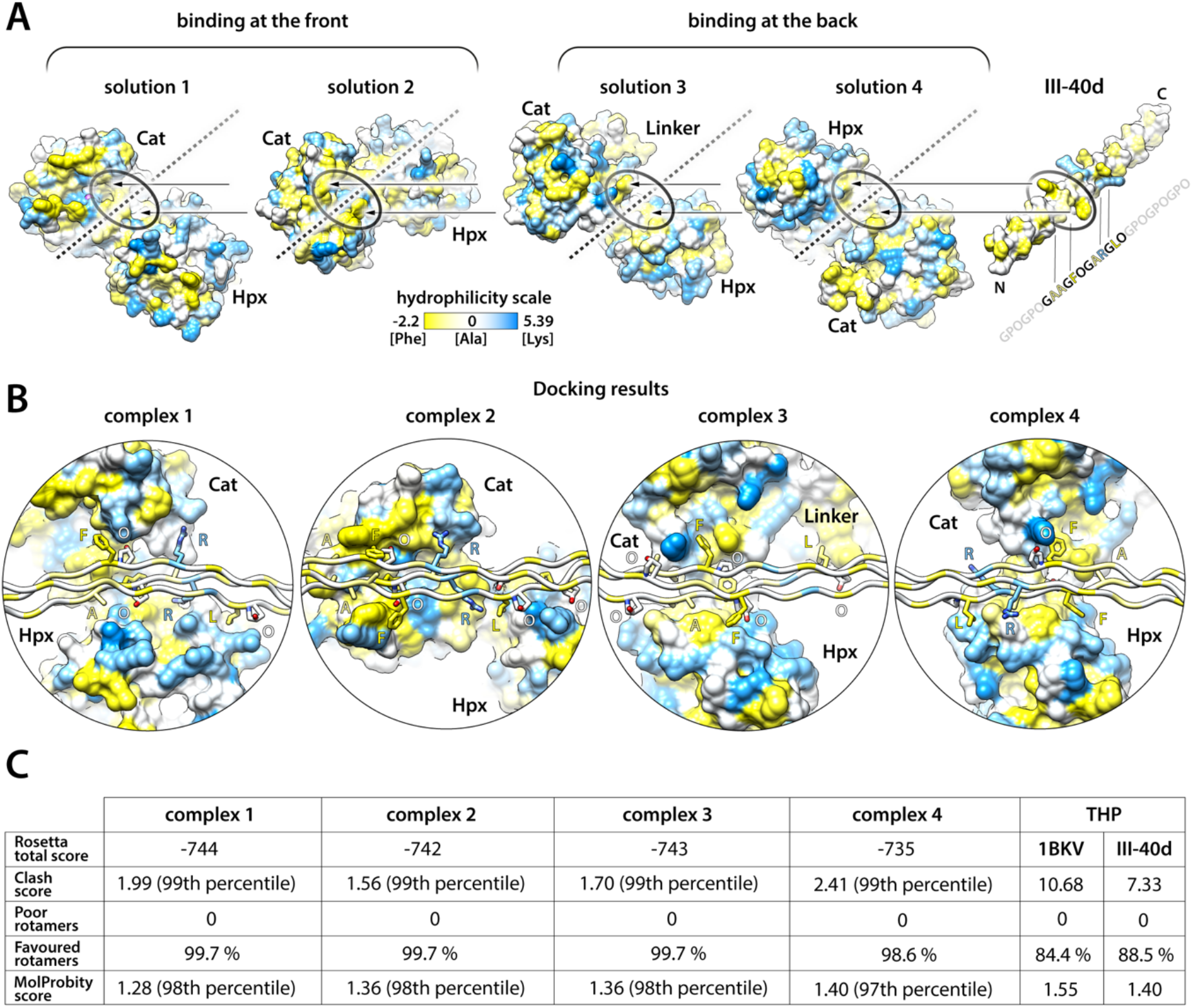
Four potential collagen binding modes of MMP-3 guided by hydrophobic interactions. **A**) Collagen II and III recognition by MMP-3 relies on the presence of Phe at position X of the collagen GXY repeat. Thus, solvent-accessible surfaces (SAS) of MMP-3 and III-40d THP models have been coloured according to hydrophilicity scale^39^: values ranging from −2.2 for Phe (most hydrophobic) to 5.39 for Lys (most hydrophilic). Four hydrophobic (yellow) patches on MMP-3 surface that can accommodate the critical GFO/A triplet in the way that would involve both Cat and Hpx domains in the binding of the collagen triple helix are indicated with oval shapes: two are at the front, and two at the back of MMP-3, according to the standard MMP-3 orientation shown in Fig. S1. The relative orientations of MMP-3 in each of the four considered modes of collagen binding are compared in relation to a fixed orientation of the III-40d THP, as indicated with broken lines representing the THP/collagen axis. For the subsequent refinement of the MMP-3-III-40d binding interface the THP has been roughly positioned within ~5 Å distance from the MMP-3 homology model in each of the 4 considered binding modes, as indicated with the arrows. **B**) Results of the local docking and interface refinement by Rosetta^40^. Models 1-4 correspond to the putative THP/collagen binding modes 1-4 shown in A. The III-40d THP is oriented horizontally for each model, demonstrating various degrees of THP bending in the modelled complexes. MMP-3 is shown as SAS (as in A) and the THP as ribbon, with side chains of the residues containing at least one atom located within 4 Å distance from MMP-3 shown as sticks and labelled with one-letter codes; colouring as in A and by heteroatom: N, navy blue; O, red. The hydrophilicity colouring algorithm works on the per residue basis and thus colours the whole residue according to the assigned value. Thus, hydrophobicity of the aliphatic stalks of the side chains such as Lys or Arg is not accounted for. **C**) Model refinement statistics table. Rosetta total score combines multiple terms of the Rosetta energy function and is expressed in arbitrary units: lowest scores correspond to most energetically favourable complexes. Clashscore, rotamers and MolProbity scores are determined using the MolProbity^41^ server. MolProbity score combines the clashscore, rotamer, and Ramachandran evaluations into a single score. For comparison, included are the relevant scores of the THP structure solved by X-ray diffraction (PDB: 1BKV) and those of the III-40d THP built on the 1BKV backbone. Rosetta refinement has reduced internal clashes and improved rotamer conformations of III-40d in the complex with MMP-3, resulting in excellent overall stereochemistry in all four complexes (>97^th^ percentile among N=27675 structures of 0-99 Å resolution).

### Complex 2 represents the putative collagen binding mode of MMP-3

Fig. 3A shows the four candidate MMP-3-THP complexes in the standard enzyme orientation. All three α-chains make contacts with MMP-3 in all the complexes (Fig. 3B), and each complex shows several hydrogen bonds between MMP-3 and the THP involving both the Cat and Hpx domain (Fig. S3). The total MMP-3-THP interface area is largest in complex 2 (1141.7 Å^2^) and smallest, but still extensive in complex 4 (825.6 Å^2^). In theory, the enzyme could be using several modes of collagen binding with variable preference for particular sites and all four candidate complexes seem equally legitimate. Without experimental data it would be impossible to rank them in terms of the likelihood of representability of the true mechanism. However, the Toolkit screening has endowed us with knowledge of residues that are favourable and disruptive for MMP-3 binding at defined THP positions (Fig. 1D). So, we set out to validate each candidate complex by substituting selected residues in the THP model III-40d for those that showed most profound effects on MMP-3 binding. To test the expected disruptive effects, we swapped Ala and Hyp at positions Y of the Triplets −1 and 0, respectively, to Arg (Fig. 3C), which cancels MMP-3(E200A) binding (Fig. 1B and D). Only complex 2 showed severe clashes between the MMP-3 model and the Arg side chains and no obvious clashes arose in complexes 1, 3 and 4. For completeness, we also tested the expected beneficial effects of swapping Leu at position X of the Triplet +2 to Arg (Fig. 1C, D and 3C). Complex 1 showed no change upon this substitution, complexes 3 and 4 showed a single additional hydrogen bond, and complex 2 showed 4 extra hydrogen bonds (Fig. 3C). It turns out that only complex 2 is completely in line with all our experimental evidence. We therefore conclude that complex 2 represents the putative mode of collagen binding by MMP-3 and that the other three candidate complexes are probably modelling artefacts (so-called decoys).

**Figure 3.**
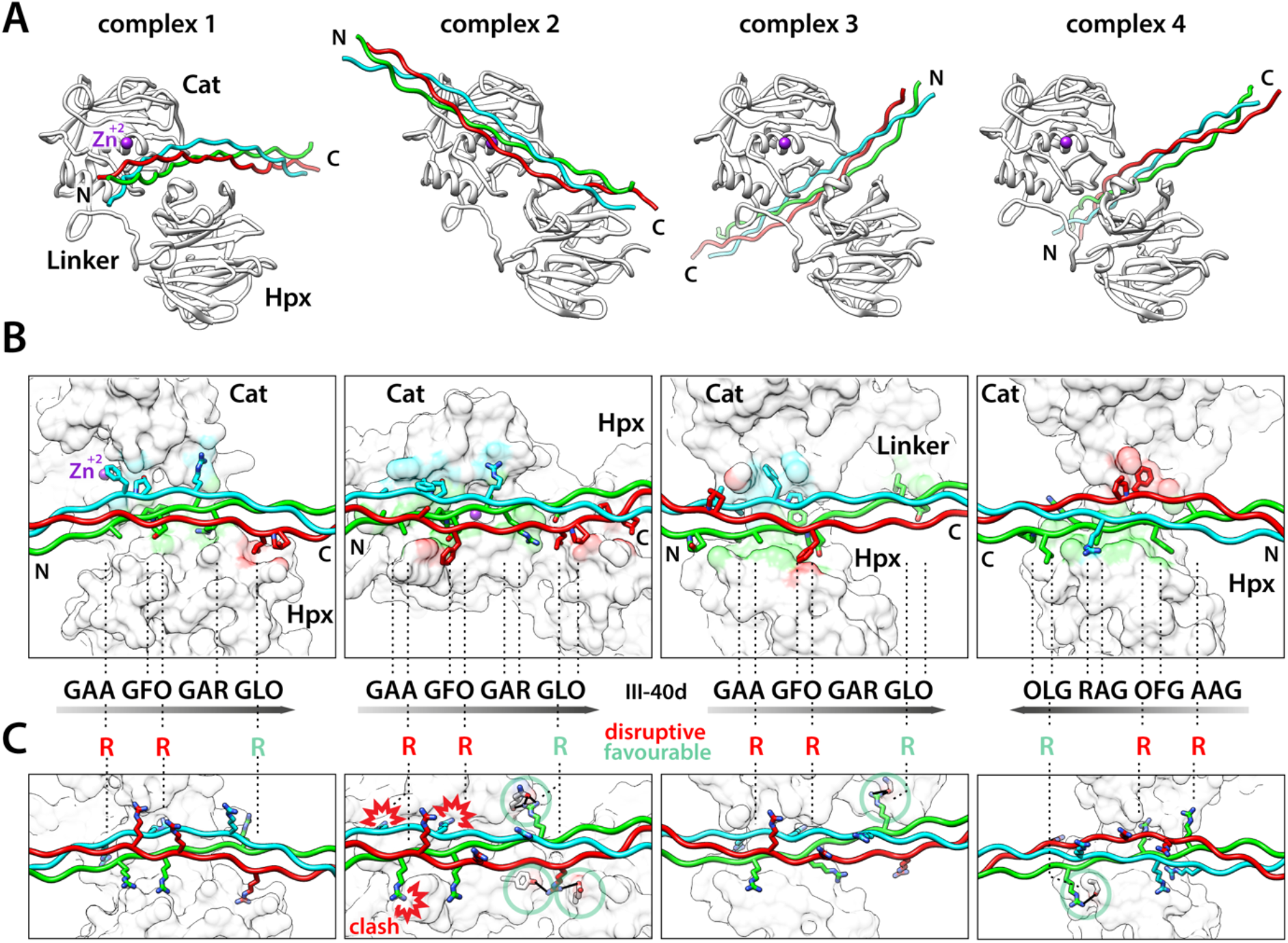
Modelling of the effects of collagen sequence variations on MMP-3 binding elects complex 2 as MMP-3’s putative collagen-binding mode. **A**) Four models of the MMP-3-III-40d complex shown as ribbons in the standard MMP-3 orientation. MMP-3 chain, white; Zn^2+^, magenta. THP chains: leading, blue; middle, green; trailing, red. **B**) Close-up view on the modelled MMP-3-III-40d interfaces. THP representation as in A, with side chains of the residues containing at least one atom located within 4 Å distance from MMP-3 shown as sticks coloured by heteroatom: N, navy blue; O, red. MMP-3 shown as solvent-accessible surface (SAS) coloured according to interfacing THP chains (4 Å distance). IDs of the staggered residues in the 4-triplet core of the III-40d THP are roughly indicated (dashed lines) together with the THP polarity (shaded arrows). In each complex all 3 chains of the THP make contacts with MMP-3. **C**) Theoretical analysis of the effects of III-40d residue substitutions on MMP-3 binding. Arg at the position Y of Triplets −1 and 0 (see Fig. 1D) causes severe clashes with MMP-3 only in model 2. Arg at the position X of Triplet 2 forms additional hydrogen bonds (black lines) with MMP-3 in models 2, 3 and 4 (green circles). MMP-3 shown as white and transparent SAS and III-40d shown as in A, with modelled Arg side chains and hydrogen-bonded MMP-3 residues represented with sticks coloured by heteroatom: N, navy blue; O, red.

### MMP-3 cleaves only unfolding collagen chains including those pre-cleaved by collagenase

The established putative mode of collagen binding by non-collagenolytic MMP-3 appears similar to that determined for collagenase MMP-1^26^, but their collagen binding specificities are completely different. We wondered what the biological role of MMP-3 binding to collagen could be. First, we investigated whether MMP-3 can directly influence collagen degradation at any stage of the process, and for that we used native tropocollagens I, II and III devoid of telopeptides to prevent fibril formation. In order to pinpoint the stage of collagen breakdown at which MMP-3 may be relevant, we needed to establish the temperature at which our collagen samples and their collagenase cleavage products (the ¾ and ¼ fragments) lose triple-helicity (T_m_, melting point). The thermal stability of tropocollagens and their fragments after digestion with MMP-1 was measured using circular dichroism (Fig. 4A). Collagen II helix was most thermostable (T_m_ = 43.3 °C) and collagen III the least (T_m_ = 40.9 °C). Likewise, the ¾ and ¼ fragments of collagen II were most thermostable (T_m_ = 39.9 °C) and those of collagen III the least (T_m_ = 36.4 °C). The triple helix is normally resistant to trypsin, which can thus be used as a reporter of collagen triple-helicity. Our trypsin digestion test revealed local thermal instability of the collagen III fold (sensitivity to trypsin) already at 35 °C (Fig. 4B), which is below the threshold of global collagen III unfolding (Fig. 4A). Both collagens I and II were resistant to trypsin at 35 °C, but the MMP-1-cleaved products of all the collagens were readily cleaved by trypsin at this temperature (Fig. 4B), which indicates that they already start to unfold at 35 °C. This is apparent in the melting curves of the collagen I and III fragments, but not in that of collagen II fragments. Consistently, collagen II fragments showed the lowest susceptibility to trypsin at 35 °C (Fig. 4B). We confirmed that at 35 °C MMP-3 was in principle unable to cleave the native triple helix, although collagen III appeared marginally cleaved approximately at the collagenase cleavage site after 8 h (Fig. 4C). This weak activity against collagen III has been reported^42^ and is consistent with the local thermal lability (breathing) of this collagen at 35°C (Fig. 4B) and with the marginal binding of MMP-3(E200A) to THP III-44 (Fig. 1A), where the collagenase cleavage site resides. To test if MMP-3 binding to collagen has any effect on collagenolysis by MMP-1, we compared the collagenolytic activity of 5 nM MMP-1 in the presence or absence of proteolytically inactive 200 nM MMP-3(E200A). The binding of MMP-3(E200A) to collagen did not interfere with collagenolysis by MMP-1 (Fig. 4D). Finally, we tested MMP-3 activity at 35 °C on the labile ¾ and ¼ fragments of collagens I, II and III generated by MMP-1. The fragments of heterotrimeric collagen I (two α1(I) chains and one α2(I) chain) were relatively readily cleaved by 200 nM MMP-3, especially the α2(I) chain, while those of homotrimeric collagens II and III were poorly cleaved (Fig. 4E), even though collagen III fragments were least thermostable (Fig. 4A). Overall MMP-3 shows gelatinase activity, as the collagen fragments that begin to melt at 35 °C (Fig. 4B) are equivalent of gelatin (denatured collagen) for the enzyme.

**Figure 4.**
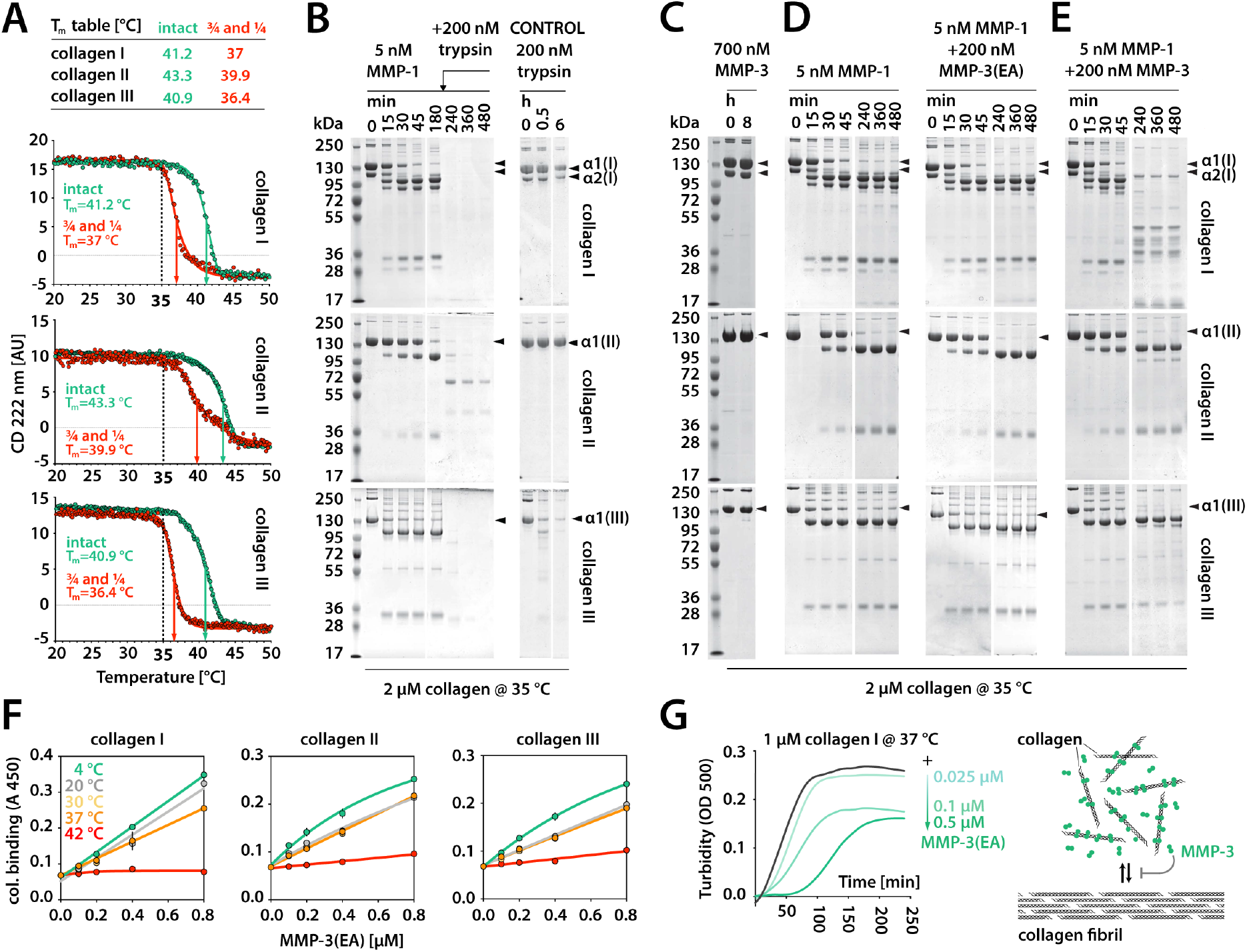
MMP-3 binding to fibrillar collagens I, II and III relies on the triple-helical fold of collagen and perturbs collagen assembly into higher order structures. **A**) Circular dichroism (CD) analysis of the melting temperatures (T_m_) of native fibrillar collagens and their MMP-1 cleavage products (¾ and ¼ fragments). **B-E**) SDS-PAGE analyses of various collagen digestion experiments. **F**) Temperature-dependent binding of biotinylated MMP-3(EA) to collagens I, II and III. The results are the average of a triplicated assay. Error bars, standard deviation (SD), smaller than the data point where not visible.

### MMP-3 affects collagen fibrillogenesis *in vitro*

To test the importance of the triple-helicity of collagen for MMP-3 binding we performed the solid phase binding assay at various temperatures using collagens I, II and III as substrates (Fig. 4F). In line with our molecular modelling experiment (Figs. 2 and 3) collagen binding to MMP-3(E200A) is highly dependent on the triple-helical structure (Fig. 4F): the lower the temperature, the higher the binding. This may explain the overall weak proteolytic activity of MMP-3 on the gelatinised (unfolding) collagenase cleavage products (the ¾ and ¼ fragments) at 35 °C (Fig. 4E). We then used collagen I sample containing telopeptides to test whether MMP-3 binding to such fibril-forming collagen may interfere with the fibril formation. As expected from the number of MMP-3 binding sites along tropocollagens II and III, MMP-3(200A) dose-dependently slowed down collagen I fibrillogenesis at 37 °C, as measured by turbidimetry at 500 nm wavelength. This suggests that one of the direct functions of MMP-3 binding to the triple helix is to regulate collagen fibrillogenesis.

## Discussion

Due to many overlapping activities and biochemical properties among MMPs and because of complex pleiotropic effects that the presence or absence of a particular MMP may invoke under various physiological or pathological circumstances, it is extremely hard to unravel the exact role of any particular MMP. To approach this question, structural insights are helpful, as every mechanistic clue is a leap in understanding of the functional potential. Here we focused on MMP-3 (stromelysin 1) which, like other MMPs, can mediate a vast number of events in the body. Its prolonged presence in tissues via immobilisation on collagen, the most abundant protein in the body, would therefore be expected to have profound consequences. The goal of this study was to gain better understanding of these potential consequences through elucidation of the structural details of MMP-3 interaction with interstitial fibril-forming collagens.

The identification of multiple MMP-3 binding sites on collagens II and III, each exhibiting different affinity to MMP-3, explains the previous finding of hardly saturable collagen binding by MMP-3^25^ also seen here (Fig. 4F). These binding sites are distinct from those previously identified with the Toolkit screening for MMP collagenases: MMP-1^26^ and MMP-13^27^. Despite homology, MMP-3 fails to effectively recognise the unique site on interstitial collagens that is recognised by these collagenases, which may at least in part explain why MMP-3 is not collagenolytic. The triple helix binding mechanism relies on different structural features in these enzymes, although for both MMP-1 and MMP-3 it is generally driven by hydrophobic interactions and involves both Cat and Hpx domains^26^.

An important difference between the collagenolytic MMP-1 and non-collagenolytic MMP-3 is that MMP-1 prefers a loose triple helix at 25-35 °C and has low affinity to both tight triple helix at 4 °C and gelatin at 42 °C, whereas MMP-3 prefers a tight triple helix at 4°C and its affinity drops with decreasing triple-helicity at increasing temperatures (Fig. 4F). This is in line with our molecular modelling experiment (Fig. 2 and 3) which shows how snuggly the triple helix fits in a valley running along the two domains of MMP-3. This putative collagen binding mode, represented by our complex 2, assumes that the main hydrophobic interaction occurs in the Cat domain, yet this domain alone is insufficient for collagen binding (Fig. 1A). This suggests that the Hpx domain plays an important role in positioning of the enzyme on the substrate. Similar cooperation between the domains is seen in MMP-1 interaction with the triple helix^26^, which shows a similar interface area between the binding partners (1290 A^2^ for MMP-1-THP vs 1141.7 Å^2^ for MMP-3-THP in complex 2). However, MMP-1 binding to the triple helix involves another hydrophobic (Leu) cluster, 9 residues upstream of the Leu cluster that binds to the Cat domain. That second Leu cluster provides a strong anchoring to the Hpx domain (exosite binding), which is predicted to be essential for triple helix unwinding^26,43^, the step required for collagenolysis^44^. There is no evidence of a similar mechanism for MMP-3, which we find to be using a very compact recognition motif of a single triplet GFO/A. This may be another reason why MMP-3 cannot unwind and cleave collagen^44^.

Our MMP-3-THP complex prediction results from incorporation of multiple lines of experimental evidence in the modelling process. Such a hybrid approach is increasingly more common in cases of proteins and protein complexes resistant to crystallisation and too small for cryo-EM structure determination. So far there are no X-ray, NMR or cryo-EM structures of full-length MMP-3, let alone a complex with THP, but this study provides key insights for a crystallisation trial of such a complex. Use of a relatively long THP, which spanned all interaction sites with MMP-1 and extended well beyond the globular shape of the enzyme proved beneficial for the crystal lattice formation in the case of the MMP-1-THP complex^26^. The same strategy could also lead to successful crystallisation of an MMP-3-THP complex, which is probably still too small (~50 kDa) for cryo-EM. Alternatively, microelectron diffraction (micro-ED) could be useful in this case, as it works even with very small ‘invisible’ crystals^45^.

Considering the biological significance of collagen binding by MMP-3 it is fascinating to discover that it can compete for binding sites with other fibrillar collagen ligands and thus regulate their functions (Fig. 5A and B). These include the following proteins screened for Toolkit binding in the past, recently reviewed^29^: i) discoidin domain receptors (DDR1 and DDR2)^46^, which control mammary tissue (DDR1), long bones (DDR2), and are potentially involved in fibrotic states, atherosclerosis and cancer; ii) von Willebrand factor (vWF)^47^, essential for platelet adhesion to damaged blood vessel walls and in blood coagulation; iii) matricellular calcium binding protein SPARC (secreted protein acidic and rich in cysteine)^48^ also known as BM40 or osteonectin, which is counter-adhesive and anti-proliferative; iv) platelet activatory receptor glycoprotein GpVI^49^; and v) osteoclast-associated receptor (OSCAR), participating in osteoclastogenesis^50^ (Fig. 5A and B). In addition to the potential competition for the shared binding site on collagen II and III, MMP-3 can cleave SPARC into 3 biologically active peptides: Z-1, which increases angiogenesis, and Z-2 and Z-3, which inhibit cell proliferation^51^.

**Figure 5.**
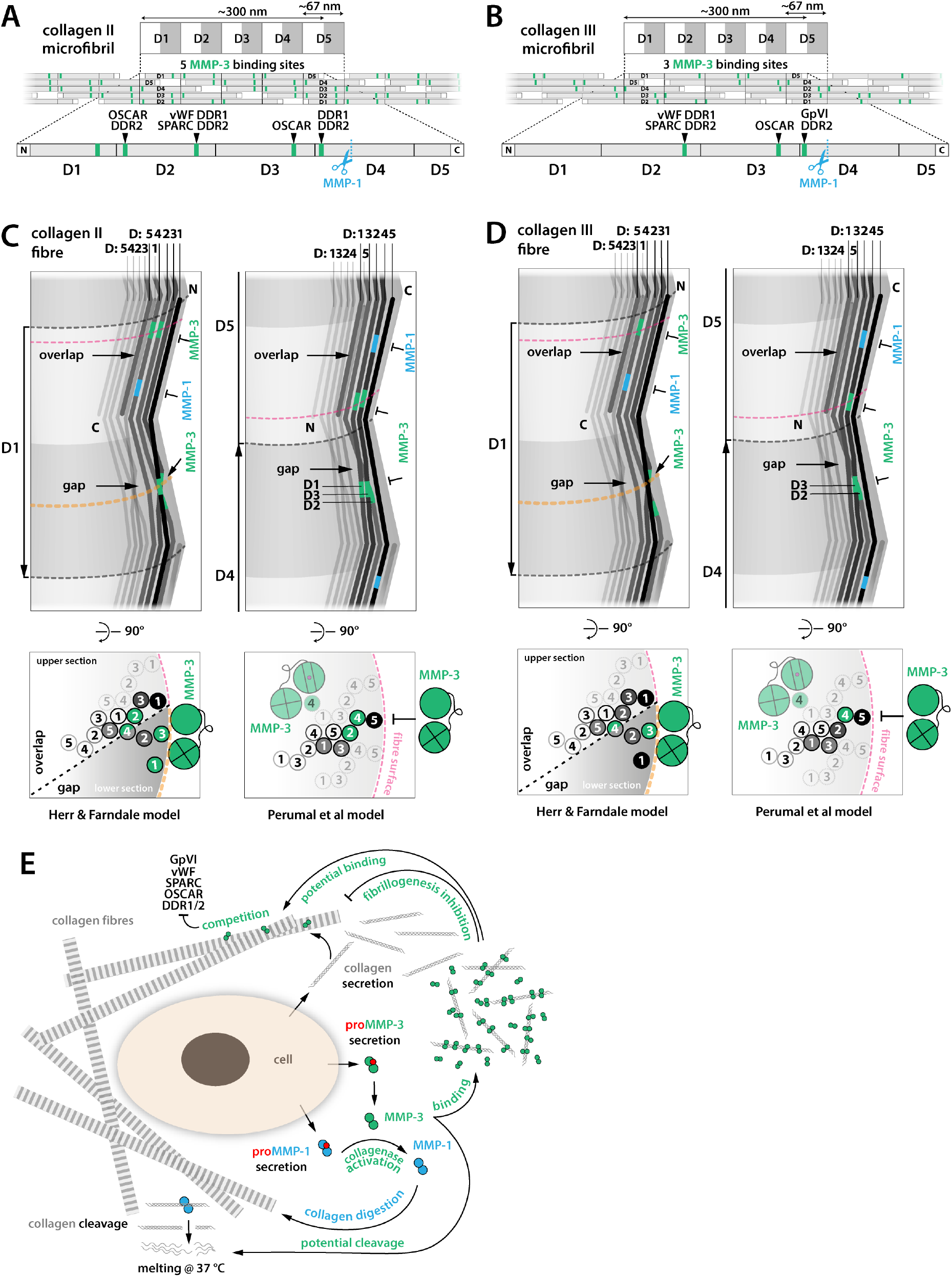
MMP-3 can regulate collagen turnover and functions of multiple collagen-binding proteins. **A and B**) Schematic microfibrillar organisation of collagen molecules forming gap and overlap regions resulting in the classical periodic D-banding pattern observed with heavy atom staining under electron microscope, due to stain accumulation in the gap regions. Each D-period contains the entire sequence of the collagen molecule distributed between 5 collagen molecules assembled with 1D stagger. Numbering of D periods in the schematics of the banding pattern is according to the D-segment order of the top collagen molecules in the schematics of the microfibrillar stagger. Non-helical telopeptides on the collagen N- and C-termini are indicated with white boxes. MMP-3 binding sites overlap with the previously mapped sites of multiple collagen binding molecules: discoidin domain receptors, DDR1 and DDR2 (GxRGQOGVMGFO); von Willebrand factor, vWF (GxRGQOGVMGFO); SPARC (secreted protein acidic and rich in cysteine) or BM-40/osteonectin; platelet activatory receptor Gp (glycoprotein) VI2; and osteoclast-associated receptor, OSCAR (GxOGPxGFxGxO). **C and D**) Schematics of 3D arrangement of collagen microfibrils in the collagen fibre according to Perumal et al and Herr & Farndale models and locations of MMP-3 binding sites (green) and MMP-1 cleavage sites (blue) in each arrangement. Collagen molecules are represented with zig-zag lines with grey levels fainting towards fibre centre. Lateral views (top panels) show interdigitation between collagen molecules in neighbouring microfibrils and their winding with respect to indentations on the surface of the collagen fibre, observed with scanning electron microscopy and atomic force microscopy. The Perumal et al model assumes that the C-termini of collagen molecules are exposed on the surface of the fibre, whereas the Herr & Farndale model assumes that these are the N-termini. Fibrillar D-period labels are next to the arrows indicating fibre polarity (N → C) and are numbered according to the order of D-segments of the outer-most, surface exposed collagen molecules (black), corresponding to fibrillar D-numbering in A and B. D-segment order of the individual collagen molecules is also indicated in the distinct staggered arrangements of each fibre model. Cross-section schematics (bottom panels) show selected collagen molecules, numbered according to their D-segment in the cross-section. All MMP-3 binding sites are buried in the idealised collagen fibre according to the Perumal et al model, preventing MMP-3 access (T symbols). Only incomplete fibrillar assemblies would enable MMP-3 binding. The Herr & Farndale model predicts exposure of several MMP-3 binding sites on the fibril surface and their potential binding by MMP-3 (orange cross-sections). **E**) Schematic illustration summarising potential functions of MMP-3 in the ECM. MMP-3 may regulate collagenolysis and fibrillogenesis by: i) potent activation of the collagenolytic MMPs (MMP-1, −8, −13); ii) potential cleavage of collagen fragments as they unwind in the body after the initial MMP-1, −8, −13 or −14 cleavage; iii) binding to newly secreted collagen molecules, preventing their fibrillation and increasing their susceptibility to collagenase. MMP-3 may also regulate function of several collagen binding proteins by competing for their collagen binding sites (as shown in A).

Schematic 2D models of collagen microfibrils, illustrating their characteristic D-periodicity, show clustering of the MMP-3 binding sites in both collagen II and III microfibrils (Fig. 5A and B). However, the actual arrangement of microfibrils in the collagen fibre is still unclear and the exposure or burial of MMP-3 binding sites – and those of many other collagen-binding proteins for that matter – may critically depend on this arrangement. Two models of collagen microfibril packing in a collagen fibre have been proposed (Fig. 5C and D), based on an intermediate resolution structure of collagen fibre^52^: one assumes exposure of tropocollagen C-termini on the fibre surface (Perumal et al model^53^), while the other assumes the exposure of the N-termini (Herr & Farndale model^54^). Each model also results in distinct interdigitation between individual tropocollagens of the constituent microfibrils (Fig. 5C and D). Only the Herr & Farndale model predicts some of the MMP-3 binding sites be exposed on the idealised, intact surface of the fibre. The Perumal et al model predicts all MMP-3 binding site be cryptic in such a fibre and MMP-3 could only bind to incomplete or damaged fibres (Fig. 5C and D). Others suggest through molecular dynamics simulations that collagen fibres are dynamic and ‘smart’, as they sample a collection of states in which cryptic sites are temporarily exposed or buried^55^. One of us has concluded that the evolution of binding sites for numerous ligands that are distributed across the tropocollagen molecule favours accessibility of collagen sequence that is not related to the D-period^29^. High-resolution fibre diffraction and/or correlative super-resolution light and electron microscopy (CLEM) may in the future use MMP-3, among other collagen binders, as a reporter protein to inform on the arrangement of collagen in the fibre.

In both models of collagen fibre assembly, a number of MMP-3 binding sites are concealed, consistent with the observed inhibition of fibrillation in the presence of MMP-3 (Fig. 4G). This suggests that MMP-3 may regulate assembly of tropocollagens into higher order structures, as shown for SPARC^48^ that shares binding sites on these collagens with MMP-3. MMP-3(E200A) added at the ratio of 0.5:1 to collagen I delayed fibrillation by ~50 min, comparably to SPARC at 10:1 molar excess over collagen^48^, which indicates that MMP-3 may be very effective in blocking or regulating collagen assembly *in vivo*. This may indirectly enhance collagenolysis by rendering newly secreted collagen molecules more available to collagenolysis (Fig. 5E). It may also play a role in limiting collagen fibril diameter similarly to decorin^56^.

Both models of collagen fibre also assume that the collagenase cleavage site is concealed in the collagen fibre (Fig. 5C and D). According to the Perumal et al model it is covered by the C-telopeptide of a neighbouring tropocollagen, whereas in the Herr & Farndale model it is buried deeper in the fibre. MMP-3, among other MMPs, has telopeptidase activity and could contribute to removal of the C-telopeptides, and thus facilitate collagenolysis by increasing the exposure of the collagenase cleavage site on the surface of the fibre. This would constitute another way in which MMP-3 could regulate collagenolysis *in vivo*.

The exposure or burial of any binding sites on collagens may be largely regulated by distinct fibrillar architectures that may not be easily reduced to any model. This includes hybridisation of multiple collagens (see further), leading to species more or less decorated with certain ligands and more or less proteinase resistant. Collagen fibrils in the body range in diameter from 10 to 500 nm^6,57^, depending on the specific tissue and its state of development. Collagen I is widespread, except in cartilaginous tissues, while collagen II is specific for cartilage^58^ and they both often co-distribute with Collagens V and XI, respectively^59–61^. The hybridisation of collagens II and XI leads to formation of ‘thin’ fibrils (16 nm)^62^, co-existing with homotypic ‘thick’ (40 nm) fibrils in the cartilage^63^. Collagen III is present in elastic tissues, mainly in blood vessels and skin and is often found at the periphery of type I collagen fibrils, especially in embryonic skin^64^. Some collagen fibrils become organised into elaborate hierarchical arrays of parallel rope-like fibres and bundles of fibres in tendons and ligaments, concentric layers in bones or orthogonal lattices in the cornea. In the latter, such an arrangement pertains to small diameter fibrils (~20 nm) and is essential for optical transparency^65^. This abundance of distinct hierarchical states of fibrillar collagens illustrate the need for various tissue-specific mechanisms regulating collagen fibril formation, where MMP-3 may play a role.

The observed multi-site collagen binding also suggests the mechanism of MMP-3 tissue retention. Decoration of interstitial collagen in the extracellular space may be a way to evade endocytosis through cell surface receptors. Several serine proteinase inhibitors, such as α_2_-Antiplasmin, α_1_-Proteinase inhibitor and α_1_-Antichymotryspin can be cleaved and inactivated by MMP-3^16^. This is important, as serine proteases like plasmin, plasma kallikrein or neutrophil elastase initiate proMMP-1 and proMMP-3 activation^66^. After such initial cleavage in the pro-domain, proMMP-3 autoactivates through autolysis and then completes activation of partially activated procollagenases. This complex network of reciprocal relations shows tight regulation of proteolytic activity on one hand, but on the other hand a potential for a positive feedback loop in case the enzyme escapes the inhibitory mechanisms. Such positive feedback may lead to rapid hyperactivation of MMP-3 along with many other MMPs, especially those downstream of MMP-3 in the activation cascade such as the collagenases. Tissue destruction in numerous diseases can be attributed to such events. For example, aberrant collagenolysis is greatly manifested during joint destruction in rheumatoid and osteoarthritis. MMP-3 is upregulated post-myocardial infarction^17^ and in a mouse model, overexpression of MMP-1 caused functional heart defects due to a reduction in collagen^67^. MMP-1 is also one of several MMPs implicated in osteoporosis^68^, since collagen I degradation by MMP-1 initiates bone resorption by activating cathepsin K producing osteoclasts^69,70^. This poses an interesting question, whether the binding of the major procollagenase activator MMP-3 to collagen matrix may constitute a reservoir of immediately bioavailable and uninhibited enzyme, ready to initiate a chain reaction when triggered by inflammation or a specific external factor. We showed that MMP-3 binding to tropocollagens I, II and III does not protect them against collagenolysis by MMP-1. According to our study, after collagen cleavage by MMP-1 or another collagenase, MMP-3 would be expected to lose affinity for the melting ¾ and ¼ fragments and dissociate from them. Once liberated from the collagen storage it could boost the collagenolytic process by activating more procollagenases and by inactivating proteinase inhibitors, which could in turn lead to even more enhancement of MMP activation through serine proteinases. As previously outlined, MMP-3 could also aid collagenolysis by removing collagen telopeptides and by inhibiting fibrillogenesis of the newly secreted tropocollagens, increasing the collagenase access to the scissile bonds. Finally, it could also contribute directly to chopping of the pre-cleaved, unfolding collagen α-chains into smaller fragments (Fig. 4E). This essentially gelatinolytic activity did not appear potent in our hands and we think it is unlikely to be of great significance *in vivo*, considering the presence of proficient gelatinases, such as MMP-2 and −9, along with other proteases that can cleave unfolded collagen α-chains in the body. Nevertheless, MMP-3 treatment of *ex vivo* cartilage explants showed generation of several collagen II fragments^71^. Crucially, these fragments do not correlate with the MMP-3 binding sites mapped in this study, suggesting that MMP-3 binding to collagen II is unrelated to its gelatin cleavage specificity.

In summary, our work explains the mechanistic differences between a prototypic collagenase MMP-1 and non-collagenolytic stromelysin MMP-3, encouraging a new perspective that closely related yet different MMPs can effectively act as stage-specific collagen chaperones, with collagenases capable of inducing and chaperoning (stabilising) the unwound state of the collagen helix in order to subsequently cleave it, and stromelysins specialising in multi-site interactions with the tight triple-helical conformation to regulate collagen fibril assembly and in turn collagenolysis. It also defines a model, likely applicable to other extracellular enzymes, where a single enzyme can directly, as well as indirectly and irrespective of its enzymatic activity, regulate aspects of ECM dynamics and activities of other extracellular proteins sharing common binding substrates. As such, the described functional potential of MMP-3 may be highly relevant in normal development, adult tissue homeostasis and diseases where dysregulated matrix turnover prevails.

## Acknowledgements

S.W.M. thanks Emeritus Professor Hideaki Nagase of Oxford University and Emeritus Professor Gillian Murphy of Cambridge University for encouragement, mentorship, many invaluable discussions and critical reading of this manuscript. We thank Dr Rob Visse of University of Oxford for purification of collagen I and for carrying out circular dichroism experiments, and Dr Shunji Hattori of Nippi Research Institute of Biomatrix in Toride, Ibaragi, Japan, for his gift of bovine collagen III. This work was supported by grants to R.W.F. from Wellcome Trust (094470/Z/10/Z) and British Heart Foundation (RG/09/003/27122).

## Author Contributions

S.W.M. designed and carried out experiments and computations, analysed data, interpreted results and wrote the manuscript; D.B. synthesised the THP Toolkit; R.W.F acquired funding, administered the project, reviewed and edited the manuscript.

## Declaration of Interests

The authors declare no competing interests.

## Methods

### Synthesis of collagen peptides (THP Toolkits)

Human collagen II and III THP Toolkits were synthesized by Fmoc (N-(9-fluorenyl)-methoxycarbonyl) chemistry as C-terminal amides on TentaGel R RAM resin using a CEM Liberty microwave-assisted automated peptide synthesizer and purified as described^72^. For exact sequences see references^46,72^. All peptides were verified by mass spectrometry and shown to adopt triple-helical conformation by polarimetry.

### Expression, refolding and purification of recombinant human MMP constructs

ProMMP-3, proMMP-3(E200A), proMMP-3Cat, MMP-3Hpx and proMMP-1 were generated according to the methods described by^44^. They were overexpressed from a pET3a vector in *E. coli* BL21 (DE3) strain (Invitrogen). Transformed cells were grown to OD600 ~ 0.4, then induced with 0.5 mM isopropyl-β-D-thiogalactopyranoside (IPTG), and harvested after 4 h. Inclusion bodies were collected by lysing the cells in 0.05 M Tris-HCl pH 8, 0.1 M NaCl, 1 mM ethylenediaminotetraacetic acid (EDTA), 0.26 mg/ml lysozyme (Sigma-Aldrich), 0.5% Triton-X100, and dissolved in 8 M Urea, 50 μM ZnCl_2_, 20 mM Tris-HCl, pH 8.6, 20 mM dithiothreitol (DTT). This was passed over a Macroprep HighQ ion-exchange column (BioRad), equilibrated in 8 M Urea, 50 μM ZnCl_2_, 20 mM Tris-HCl, pH 8.6, 1 mM DTT and eluted with a linear salt gradient (0-0.5 M NaCl). The fractions were run on SDS-PAGE and the relevant protein fractions were pooled, diluted with 50 mM Tris-HCl, pH 8.6, 6 M Urea, 1 mM DTT, 150 mM NaCl, 5 mM CaCl_2_, 100 μM ZnCl_2_, 0.02 % NaN_3_ to A280 < 0.3, supplemented with 20 mM cystamine and refolded by dialyses at 4 °C against 4 volumes of renaturation buffer (50 mM Tris-HCl, pH 8.6, 150 mM NaCl, 5 mM CaCl_2_, 100 μM ZnCl_2_, 5 mM β-mercaptoethanol, 1 mM 2-hydroxyethyl disulphide, 0.02 % NaN_3_) for 24 h, and then 10 volumes of the same buffer for another 24 h, then against 10 volumes of the same buffer without β-mercaptoethanol for 24h, and finally against 4 volumes of 50 mM Tris-HCl, pH 8.6, 5 mM CaCl_2_, 50 μM ZnCl_2_, 0.02 % NaN_3_, for 24 h. Refolded protein was purified using Green A affinity column (Amicon), equilibrated with 50 mM Tris-HCl pH 7.5, 75 mM NaCl, 5 mM CaCl_2_, 0.02 % NaN_3_, and eluted with linear salt gradient (0-1 M NaCl). ProMMPs were activated with MMP-3Cat in 50:1 molar ratio and 1 mM p-aminophenyl mercuric acetate (APMA) (ICN Biochemicals) in TNC buffer (50 mM Tris-HCl pH 7.5, 150 mM NaCl, 10 mM CaCl_2_, 0.02 % NaN_3_) for 60-120 min at 37 °C. The mature forms were purified by Sephacryl S200 gel filtration (GE Healthcare) in TNC buffer.

### Protein biotinylation

To avoid the presence of primary amines, the TNC buffer in which proteins were stored was exchanged using Sephadex G-25M PD-10 desalting gravity columns (GE Healthcare) into 50 mM N-Cyclohexyl-2-aminoethanesulfonic acid (CHES) buffer pH 8.8, supplemented with 200 mM NaCl and 10 mM CaCl2. Then, 10 mM EZ-Link Sulfo-NHS-LC-Biotin (Thermo Fisher Scientific) solution in distilled water was added at 1:2 protein-biotin molar ratio and incubated for 1 h at room temperature. Proteins were next passed over another PD-10 column equilibrated in TNC buffer to remove excess biotin.

### Acquisition of collagens I, II and III

Collagen I was extracted from Guinea pig dermis. The skin was extensively scraped, cut into pieces, washed with saline and extracted with 0.5 M acetic acid. A part of the prep was treated with pepsin for removal of the non-triple-helical telopeptides. Pepsin was added to 1/50 of the total wet weight for 24 h at 4 °C, and collagen was purified as described in the reference^73^. The pepsin-treated sample is less prone to fibrillogenesis. The non-pepsin-treated sample was stored as the fibrillogenesis-competent sample. Final collagen yield and concentration in both samples was determined after freeze-drying. Collagen II was a pepsin-digested guanidine-HCl-extract from a bovine joint cartilage purchased from Sigma-Aldrich/Merck. Bovine Collagen III was a gift from Dr. Shunji Hattori of Nippi Research Institute of Biomatrix in Toride, Ibaragi, Japan.

### THP Toolkits and collagen binding assays

For the Toolkits screening, Costar^®^ High Binding 96-well microtiter plates (Corning, UK) were coated with a 5 μg/ml THP solution in 10 mM acetic acid, incubated overnight at 4 °C. They were then washed with TNC buffer containing 0.05 % Tween 20 (TNC-T) and blocked with 3% bovine serum albumin (Sigma) in TNC-T buffer. Biotinylated proteins at 1 μM concentration were added and incubated 1-2 h at room temperature. Plates were developed using streptavidin-horseradish peroxidase conjugate (R&D, UK) and 3,3’,5,5’-tetramethylbenzidine 2-Component Microwell Peroxidase Substrate KitTM (KPL, UK) for a fixed time.

For collagen binding assay, the Costar plates were coated with 50 μl of 20 μg/ml collagen I, II or III in TNC buffer, incubated overnight at room temperature, washed and blocked as described above. Biotinylated proteins at increasing concentrations were added in TNC buffer and incubated for 2 h at 4-40 °C. The wells were then washed in TNC-T buffer at the temperature of incubation and subsequently fixed with 3 % p-formaldehyde for 30 min. Plates were developed as in the Toolkits binding assay. All assays were carried out in a triplicate and compared analyses were always developed simultaneously.

### Thermostability of collagens I, II and III and their MMP-1 cleavage products

Melting curves of collagens and their ¾ and ¼ fragments generated by collagenase were determined using circular dichroism (CD). Proteins were diluted to 10 μM concentration in TNC buffer and their thermal transitions were monitored by ellipticity (Θ) change at 222 nm across 0.1 cm pathlength in a Jasco 815 CD instrument. Temperature was increased at 0.1°C/min.

### Collagenolysis assays

MMPs or trypsin were incubated with collagen in TNC buffer at indicated concentrations. Reactions were stopped at different time-points by the addition of an equal volume of the reducing ammediol loading mix [42 mM ammediol-HCl pH7.5, 0.01 % (w/v) NaN_3_, 2 % (w/v), sodium dodecyl sulfate (SDS), 50 % (w/v) glycerol, 1 % β-mercaptoethanol and a few grains of bromophenol blue] containing 20 mM EDTA and 10 mM phenylmethanesulfonyl fluoride (PMSF). The cleavage products were analysed by SDS-PAGE using a modification of the ammediol-glycine gel and buffer system with 7.5 % total acrylamide and the gels were stained with Coomassie Brilliant Blue R-250.

### Turbidimetric collagen I fibrillogenesis assay

Collagen fibrillation can be monitored by the extent of light scattering or turbidity. Fibrillogenesis competent collagen I (non-pepsin-digested) at the concentration of 1 μM was mixed with MMP-3(E200A) at indicated concentrations in 20 mM Hepes pH7.4, 150 mM NaCl, 2 mM CaCl2 in an Eppendorf UVette (220-1600 nm). The increasing sample turbidity was measured at 500 nm wavelength over for 4 h at 37 °C with Cary 3 UV/VIS spectrophotometer equipped with a temperature control unit (Varian).

### Homology modelling and docking

ProMMP-3 and mature MMP-3 models were obtained by homology modelling using Modeller 9.22 version^35^ via web service with default settings. The THP III-40d model for docking to MMP-3 model was obtained through replacement of residues within the 1BKV structure^30^ using Coot^31^ and UCSF Chimera^74^. The most common side chain conformations according to Dunbrack rotamer library^75^ were selected for this starting model and then refined during the Rosetta-assisted docking. The docking involved manual positioning of the THP model III-40d within 2-6 Å distance from the MMP-3 model in 4 indicated orientations, to decrease the global conformational search space and improve the efficiency of the subsequent interface optimisation with Rosetta *relax* protocol^40^. The *relax* algorithm relieved clashes in the assembly and move it to the nearest local minimum in the Rosetta energy function.

### Structure visualisation

Structural analyses and presentations were done using PDBePISA (http://www.ebi.ac.uk/msd-srv/prot_int/cgi-bin/piserver) and UCSF Chimera^74^.

## References

1. Brown, J. C. & Timpl, R. The collagen superfamily. Int. Arch. Allergy Immunol. 107, 484–490 (1995).

2. Kadler, K. Extracellular matrix 1: Fibril-forming collagens. Protein Profile 2, 491–619 (1995).

3. Engel, J. & Prockop, D. J. The zipper-like folding of collagen triple helices and the effects of mutations that disrupt the zipper. Annu Rev Biophys Biophys Chem 20, 137–152 (1991).

4. Kadler, K. E., Holmes, D. F., Trotter, J. A. & Chapman, J. A. Collagen fibril formation. Biochem. J. 316 (Pt 1), 1–11 (1996).

5. Ottani, V., Raspanti, M. & Ruggeri, A. Collagen structure and functional implications. Micron 32, 251–260 (2001).

6. Fleischmajer, R., Perlish, J. S., Timpl, R. & Olsen, B. R. Procollagen intermediates during tendon fibrillogenesis. J. Histochem. Cytochem. 36, 1425–1432 (1988).

7. Craig, A. S., Birtles, M. J., Conway, J. F. & Parry, D. A. An estimate of the mean length of collagen fibrils in rat tail-tendon as a function of age. Connect. Tissue Res. 19, 51–62 (1989).

8. Woessner, J. F. Matrix metalloproteinases and their inhibitors in connective tissue remodeling. FASEB J. 5, 2145–2154 (1991).

9. Manon-Jensen, T., Kjeld, N. G. & Karsdal, M. A. Collagen-mediated hemostasis. J. Thromb. Haemost. 14, 438–448 (2016).

10. Nagase, H. & Woessner, J. F. Matrix metalloproteinases. J. Biol. Chem. 274, 21491–21494 (1999).

11. Nagase, H., Visse, R. & Murphy, G. Structure and function of matrix metalloproteinases and TIMPs. Cardiovasc. Res. 69, 562–573 (2006).

12. Sternlicht, M. D. & Werb, Z. How matrix metalloproteinases regulate cell behavior. Annu. Rev. Cell Dev. Biol. 17, 463–516 (2001).

13. Maskos, K. Crystal structures of MMPs in complex with physiological and pharmacological inhibitors. Biochimie 87, 249–263 (2005).

14. Suzuki, K., Enghild, J. J., Morodomi, T., Salvesen, G. & Nagase, H. Mechanisms of activation of tissue procollagenase by matrix metalloproteinase 3 (stromelysin). Biochemistry 29, 10261–10270 (1990).

15. Knäuper, V., López-Otin, C., Smith, B., Knight, G. & Murphy, G. Biochemical characterization of human collagenase-3. J. Biol. Chem. 271, 1544–1550 (1996).

16. Nagase, H. Matrix Metalloproteinase 3/Stromelysin 1. in Handbook of Proteolytic Enzymes 763–774 (Elsevier, 2013). doi:10.1016/B978-0-12-382219-2.00158-7

17. Kelly, D. et al. Circulating stromelysin-1 (MMP-3): a novel predictor of LV dysfunction, remodelling and all-cause mortality after acute myocardial infarction. Eur. J. Heart Fail. 10, 133–139 (2008).

18. Lochter, A. et al. Matrix metalloproteinase stromelysin-1 triggers a cascade of molecular alterations that leads to stable epithelial-to-mesenchymal conversion and a premalignant phenotype in mammary epithelial cells. J. Cell Biol. 139, 1861–1872 (1997).

19. Posthumus, M. D., Limburg, P. C., Westra, J., van Leeuwen, M. A. & van Rijswijk, M. H. Serum matrix metalloproteinase 3 in early rheumatoid arthritis is correlated with disease activity and radiological progression. J. Rheumatol. 27, 2761–2768 (2000).

20. Yoshihara, Y. et al. Matrix metalloproteinases and tissue inhibitors of metalloproteinases in synovial fluids from patients with rheumatoid arthritis or osteoarthritis. Ann. Rheum. Dis. 59, 455–461 (2000).

21. Mudgett, J. S. et al. Susceptibility of stromelysin 1-deficient mice to collagen-induced arthritis and cartilage destruction. Arthritis Rheum. 41, 110–121 (1998).

22. Blom, A. B. et al. Crucial role of macrophages in matrix metalloproteinase-mediated cartilage destruction during experimental osteoarthritis: involvement of matrix metalloproteinase 3. Arthritis Rheum. 56, 147–157 (2007).

23. Clements, K. M. et al. Gene deletion of either interleukin-1beta, interleukin-1beta-converting enzyme, inducible nitric oxide synthase, or stromelysin 1 accelerates the development of knee osteoarthritis in mice after surgical transection of the medial collateral ligament and partial medial meniscectomy. Arthritis Rheum. 48, 3452–3463 (2003).

24. Okada, Y. et al. Localization of matrix metalloproteinase 3 (stromelysin) in osteoarthritic cartilage and synovium. Lab. Invest. 66, 680–690 (1992).

25. Murphy, G. et al. The role of the C-terminal domain in collagenase and stromelysin specificity. J. Biol. Chem. 267, 9612–9618 (1992).

26. Manka, S. W. et al. Structural insights into triple-helical collagen cleavage by matrix metalloproteinase 1. Proc. Natl. Acad. Sci. U.S.A. 109, 12461–12466 (2012).

27. Howes, J.-M. et al. The recognition of collagen and triple-helical toolkit peptides by MMP-13: sequence specificity for binding and cleavage. J. Biol. Chem. 289, 24091–24101 (2014).

28. Farndale, R. W. et al. Cell-collagen interactions: the use of peptide Toolkits to investigate collagen-receptor interactions. Biochem. Soc. Trans. 36, 241–250 (2008).

29. Farndale, R. W. Collagen-binding proteins: insights from the Collagen Toolkits. Essays Biochem. (2019). doi:10.1042/EBC20180070

30. Kramer, R. Z., Bella, J., Mayville, P., Brodsky, B. & Berman, H. M. Sequence dependent conformational variations of collagen triple-helical structure. Nat. Struct. Biol. 6, 454–457 (1999).

31. Emsley, P. & Cowtan, K. Coot: model-building tools for molecular graphics. Acta Crystallogr. D Biol. Crystallogr. 60, 2126–2132 (2004).

32. Jozic, D. et al. X-ray structure of human proMMP-1: new insights into procollagenase activation and collagen binding. J. Biol. Chem. 280, 9578–9585 (2005).

33. Iyer, S., Visse, R., Nagase, H. & Acharya, K. R. Crystal structure of an active form of human MMP-1. J. Mol. Biol. 362, 78–88 (2006).

34. Stura, E. A., Visse, R., Cuniasse, P., Dive, V. & Nagase, H. Crystal structure of full-length human collagenase 3 (MMP-13) with peptides in the active site defines exosites in the catalytic domain. FASEB J. 27, 4395–4405 (2013).

35. Sali, A. & Blundell, T. L. Comparative protein modelling by satisfaction of spatial restraints. J. Mol. Biol. 234, 779–815 (1993).

36. John, B. & Sali, A. Comparative protein structure modeling by iterative alignment, model building and model assessment. Nucleic Acids Res. 31, 3982–3992 (2003).

37. Melo, F., Sánchez, R. & Sali, A. Statistical potentials for fold assessment. Protein Sci. 11, 430–448 (2002).

38. Shen, M.-Y. & Sali, A. Statistical potential for assessment and prediction of protein structures. Protein Sci. 15, 2507–2524 (2006).

39. Moon, C. P. & Fleming, K. G. Side-chain hydrophobicity scale derived from transmembrane protein folding into lipid bilayers. Proc. Natl. Acad. Sci. U.S.A. 108, 10174–10177 (2011).

40. Nivón, L. G., Moretti, R. & Baker, D. A Pareto-optimal refinement method for protein design scaffolds. PLoS ONE 8, e59004 (2013).

41. Williams, C. J. et al. MolProbity: More and better reference data for improved all-atom structure validation: PROTEIN SCIENCE.ORG. Protein Science 27, 293–315 (2018).

42. Gunja-Smith, Z., Nagase, H. & Woessner, J. F. Purification of the neutral proteoglycan-degrading metalloproteinase from human articular cartilage tissue and its identification as stromelysin matrix metalloproteinase-3. Biochem. J. 258, 115–119 (1989).

43. Robichaud, T. K., Steffensen, B. & Fields, G. B. Exosite interactions impact matrix metalloproteinase collagen specificities. J. Biol. Chem. 286, 37535–37542 (2011).

44. Chung, L. et al. Collagenase unwinds triple-helical collagen prior to peptide bond hydrolysis. EMBO J. 23, 3020–3030 (2004).

45. Nannenga, B. L. & Gonen, T. MicroED opens a new era for biological structure determination. Curr. Opin. Struct. Biol. 40, 128–135 (2016).

46. Konitsiotis, A. D. et al. Characterization of high affinity binding motifs for the discoidin domain receptor DDR2 in collagen. J. Biol. Chem. 283, 6861–6868 (2008).

47. Lisman, T. et al. A single high-affinity binding site for von Willebrand factor in collagen III, identified using synthetic triple-helical peptides. Blood 108, 3753–3756 (2006).

48. Giudici, C. et al. Mapping of SPARC/BM-40/osteonectin-binding sites on fibrillar collagens. J. Biol. Chem. 283, 19551–19560 (2008).

49. Jarvis, G. E. et al. Identification of a major GpVI-binding locus in human type III collagen. Blood 111, 4986–4996 (2008).

50. Barrow, A. D. et al. OSCAR is a collagen receptor that costimulates osteoclastogenesis in DAP12-deficient humans and mice. J. Clin. Invest. 121, 3505–3516 (2011).

51. Sage, E. H. et al. Cleavage of the matricellular protein SPARC by matrix metalloproteinase 3 produces polypeptides that influence angiogenesis. J. Biol. Chem. 278, 37849–37857 (2003).

52. Orgel, J. P. R. O., Irving, T. C., Miller, A. & Wess, T. J. Microfibrillar structure of type I collagen in situ. Proc. Natl. Acad. Sci. U.S.A. 103, 9001–9005 (2006).

53. Perumal, S., Antipova, O. & Orgel, J. P. R. O. Collagen fibril architecture, domain organization, and triple-helical conformation govern its proteolysis. Proc. Natl. Acad. Sci. U.S.A. 105, 2824–2829 (2008).

54. Herr, A. B. & Farndale, R. W. Structural insights into the interactions between platelet receptors and fibrillar collagen. J. Biol. Chem. 284, 19781–19785 (2009).

55. Zhu, J., Hoop, C. L., Case, D. A. & Baum, J. Cryptic binding sites become accessible through surface reconstruction of the type I collagen fibril. Sci Rep 8, 16646 (2018).

56. Orgel, J. P. R. O., Eid, A., Antipova, O., Bella, J. & Scott, J. E. Decorin core protein (decoron) shape complements collagen fibril surface structure and mediates its binding. PLoS ONE 4, e7028 (2009).

57. Flint, M. H., Craig, A. S., Reilly, H. C., Gillard, G. C. & Parry, D. A. Collagen fibril diameters and glycosaminoglycan content of skins--indices of tissue maturity and function. Connect. Tissue Res. 13, 69–81 (1984).

58. Canty, E. G. & Kadler, K. E. Procollagen trafficking, processing and fibrillogenesis. J. Cell. Sci. 118, 1341–1353 (2005).

59. Kypreos, K. E., Birk, D., Trinkaus-Randall, V., Hartmann, D. J. & Sonenshein, G. E. Type V collagen regulates the assembly of collagen fibrils in cultures of bovine vascular smooth muscle cells. J. Cell. Biochem. 80, 146–155 (2000).

60. Eyre, D. Collagen of articular cartilage. Arthritis Res. 4, 30–35 (2002).

61. Wenstrup, R. J. et al. Type V collagen controls the initiation of collagen fibril assembly. J. Biol. Chem. 279, 53331–53337 (2004).

62. Keene, D. R., Oxford, J. T. & Morris, N. P. Ultrastructural localization of collagen types II, IX, and XI in the growth plate of human rib and fetal bovine epiphyseal cartilage: type XI collagen is restricted to thin fibrils. J. Histochem. Cytochem. 43, 967–979 (1995).

63. Kadler, K. E., Hill, A. & Canty-Laird, E. G. Collagen fibrillogenesis: fibronectin, integrins, and minor collagens as organizers and nucleators. Curr. Opin. Cell Biol. 20, 495–501 (2008).

64. Keene, D. R., Sakai, L. Y., Bächinger, H. P. & Burgeson, R. E. Type III collagen can be present on banded collagen fibrils regardless of fibril diameter. J. Cell Biol. 105, 2393–2402 (1987).

65. Hulmes, D. J. S. Building collagen molecules, fibrils, and suprafibrillar structures. J. Struct. Biol. 137, 2–10 (2002).

66. Nagase, H., Enghild, J. J., Suzuki, K. & Salvesen, G. Stepwise activation mechanisms of the precursor of matrix metalloproteinase 3 (stromelysin) by proteinases and (4-aminophenyl)mercuric acetate. Biochemistry 29, 5783–5789 (1990).

67. Kim, H. E. et al. Disruption of the myocardial extracellular matrix leads to cardiac dysfunction. J. Clin. Invest. 106, 857–866 (2000).

68. Andersen, T. L. et al. A scrutiny of matrix metalloproteinases in osteoclasts: evidence for heterogeneity and for the presence of MMPs synthesized by other cells. Bone 35, 1107–1119 (2004).

69. Sakamoto, S. & Sakamoto, M. Degradative processes of connective tissue proteins with special emphasis on collagenolysis and bone resorption. Mol. Aspects Med. 10, 299–428 (1988).

70. Holliday, L. S. et al. Initiation of osteoclast bone resorption by interstitial collagenase. J. Biol. Chem. 272, 22053–22058 (1997).

71. Zhen, E. Y. et al. Characterization of metalloprotease cleavage products of human articular cartilage. Arthritis Rheum. 58, 2420–2431 (2008).

72. Raynal, N. et al. Use of synthetic peptides to locate novel integrin alpha2beta1-binding motifs in human collagen III. J. Biol. Chem. 281, 3821–3831 (2006).

73. Miller, E. J. & Rhodes, R. K. Preparation and characterization of the different types of collagen. Meth. Enzymol. 82 Pt A, 33–64 (1982).

74. Pettersen, E. F. et al. UCSF Chimera--a visualization system for exploratory research and analysis. J Comput Chem 25, 1605–1612 (2004).

75. Dunbrack, R. L. Rotamer libraries in the 21st century. Curr. Opin. Struct. Biol. 12, 431–440 (2002).

