## Supplementary Information for "Structural studies of MMP-3 interaction with triple-helical collagen introduce the enzyme’s new roles in tissue remodelling"

### Supplementary figures

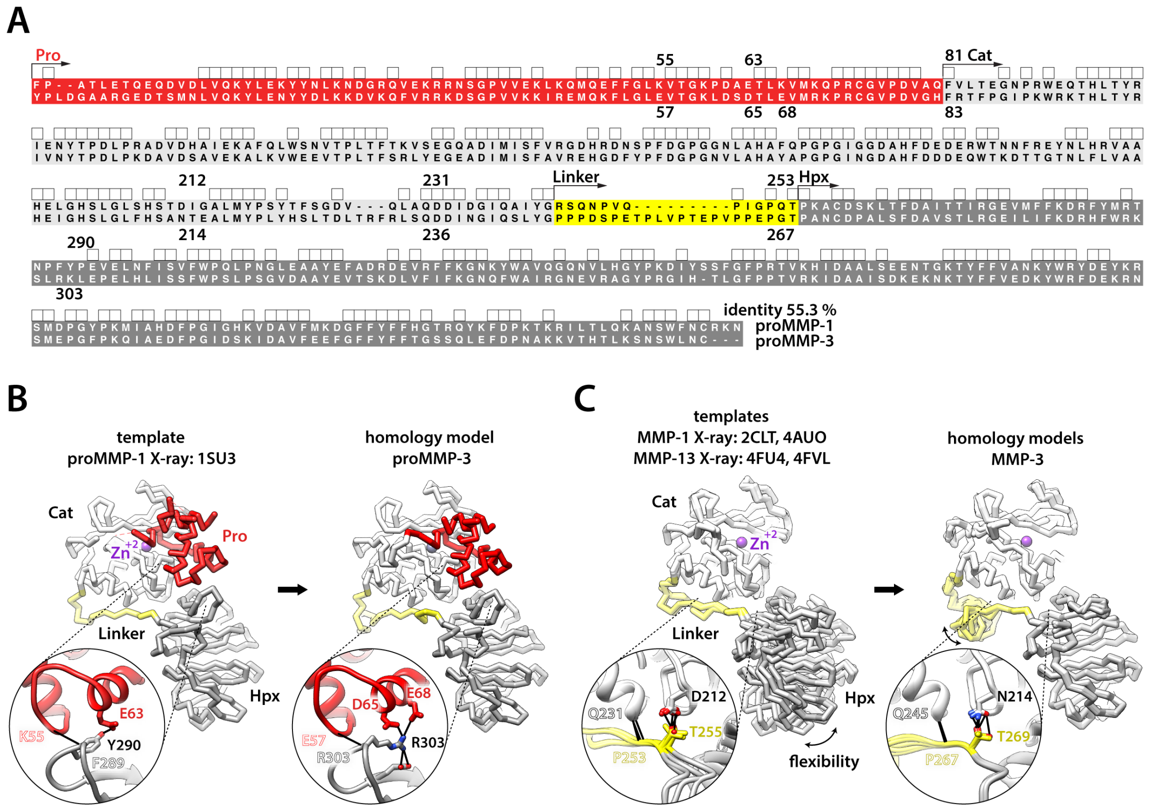


**Figure S1. Homology modelling of full-length MMP-3 in the pro- and activated form.**
**A**) Alignment of proMMP-1 and proMMP-3 sequences. Pro-domains, red; catalytic (Cat) domains, light grey; linkers, yellow; hemopexin domains (Hpx), dark grey. Selected, functionally important residues (see panels B and C) are numbered. Regions of 100 % conservation (sequence identity) are indicated with square symbols. MMP-3 linker is 9-residues shorter than that of MMP-1 (dashes). The overall sequence identity between proMMP-1 and -3 by Clustal X amounts to 55.3 %. **B**) Homology modelling by Modeller of proMMP-3, based on the only available proMMP-1 crystal structure (PDB: 1SU3; backbone representations in the standard *frontal* orientation, facing the catalytic site cleft that contains the catalytic $Zn^{2+}$ ion (magenta), with domain colouring as in A. Close-up views show the equivalent Pro-Hpx contacts (hydrogen bonds, black lines) contributing to the relative orientation of the Cat and Hpx domains in each proMMP structure/model; ribbon representation with selected residue side chains shown as sticks coloured by heteroatom (N, navy blue; O, red). Backbone contact labels, silhouette font; side-chain contact labels, solid font. **C**) Homology modelling by Modeller of activated MMP-3, based on crystal structures of full-length MMP-1 (PDBs: 2CLT and 4AUO) and MMP-13 (PDBs: 4FU4, 4FVL). Shown are the 4 template structures and the 5 resulting MMP-3 models, superposed on their respective Cat domains; representation, view and colouring as in B. The superposition reveals inter-domain flexibility via linker (hinge region) in the activated MMP reference structures (templates) and a local conformational flexibility of the unstructured linker region

in the target MMP-3 models, indicated by bent two-sided arrows. Close-up views show the equivalent linker-ordering regions along the Cat domain, focusing on selected relevant hydrogen bonds (black lines); ribbon representation with selected residue side chains shown as sticks coloured by heteroatom (N, navy blue; O, red). Backbone contact labels, silhouette font; side-chain contact labels, solid font. The models show identical overall domain folds to the templates and the relative arrangement of the domains is preserved by analogous, however distinct interactions between them (B and C, insets). Only the flexible and divergent linker, 9 aa longer in MMP-3 than in MMP-1 (A), locally samples various conformations, where it is not ordered by the stabilising interactions with the Cat domain (C).

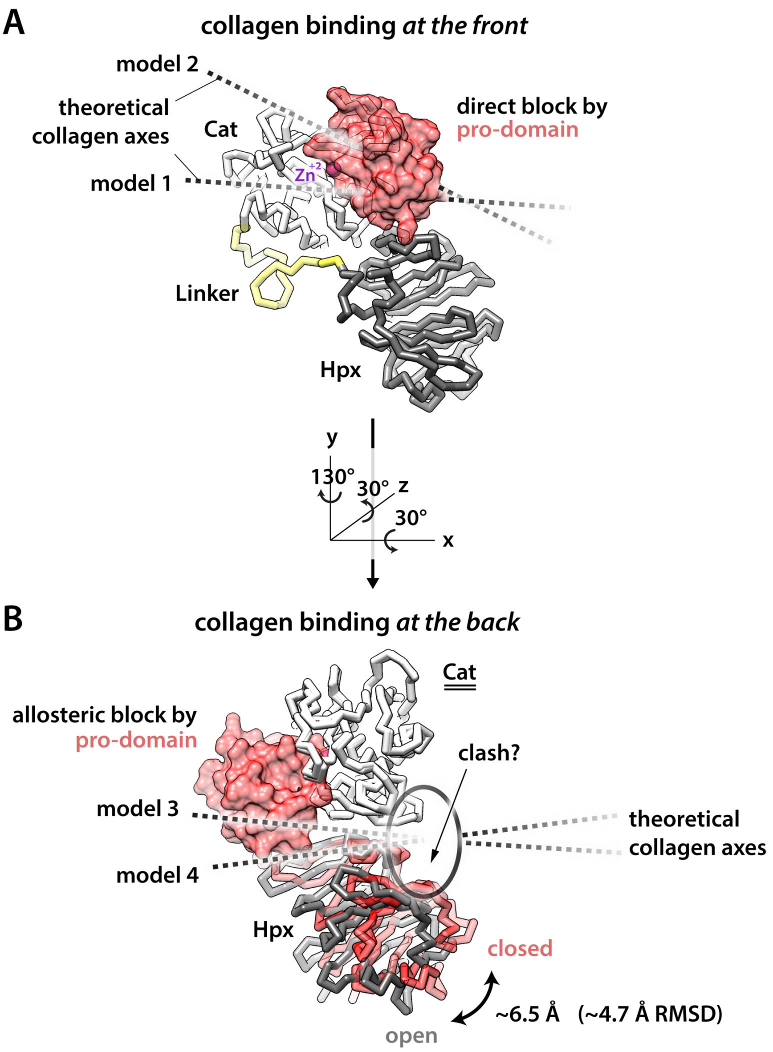


**Figure S2. Models of direct and indirect collagen binding inhibition by the pro-domain of proMMP-3.**

**A**) Direct steric hindrance would prevent collagen binding according to models 1 and 2

**B**) Pro-domain-induced change in the relative positions of the Cat and Hpx domains could indirectly prevent collagen binding according to models 3 and 4. Homology models of proMMP-3 (red) and activated MMP-3 (grey) generated using proMMP-1 and MMP-1/13 templates (Figs. 2 and S1) are superimposed on the Cat domains (parallel lines) to demonstrate the potential widening of the cleft between the Cat and Hpx domains upon proMMP-3 activation. The closed conformation may be unable to accommodate collagen molecule in its binding site (clashing with the oval shape). Shown are maximal local displacement value and RMSD (root means square deviation) between the Hpx domains in the two models.

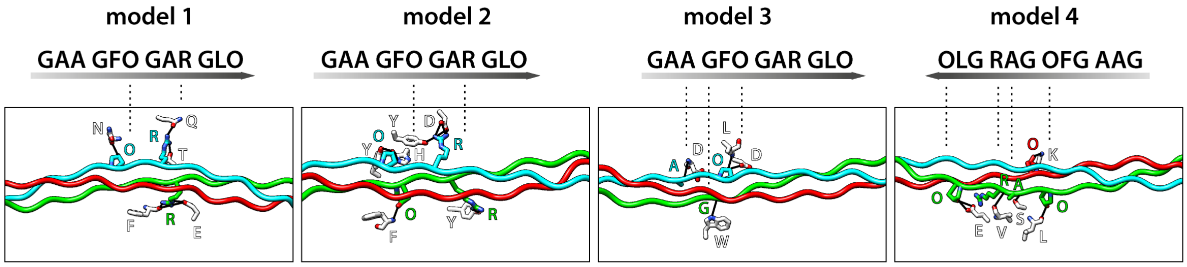


**Figure S3. Hydrogen bonding networks between MMP-3 and the THP model III-40d in the modelled complexes.**

Hydrogen bonds are shown with black lines. THP representation as in Fig. 4A, with hydrogen bonding THP and MMP-3 residues shown as sticks coloured by heteroatom: N, navy blue; O, red. IDs of the hydrogen bonding residues in the 4-triplet core of the III-40d THP are roughly indicated (dashed lines) together with the THP polarity (shaded arrows indicate N-term → C-term direction).
